# Anthropogenic subsidies reshape Ruby-throated Hummingbird (*Archilochus colubris*) interactions across spatial and temporal contexts: evidence from community-collected data

**DOI:** 10.64898/2026.07.03.733347

**Authors:** Desirée L. Narango, Amber Jones, Ryan Rebozo, Pablo Sosa-Negrón, Michael T. Hallworth

## Abstract

1. Ruby-throated hummingbirds (*Archilochus colubris*) exhibit flower preferences and readily visit human subsidies like feeders. However, foraging behavior and plant-animal interactions may vary across spatial, temporal, and landscape gradients. Despite the popularity and ubiquity of Ruby-throated Hummingbirds in the eastern U.S., there has never been a quantitative assessment of their flower preferences or feeder use across broad scales.
2. We investigated how feeder visitation, flower visitation, and flower trait preferences vary by latitude, season, and land use in the northeastern United States using over 6 million occurrence records of Ruby-throated Hummingbirds and flowering plants, >2,100 annotated hummingbird-flower interactions, and >2,700 feeder visit occurrences.
3. We found that hummingbird feeder use declined over the year, increased with latitude, and was higher in developed landscapes. Flower visitation increased over the year across all latitudes, with higher visitation in developed landscapes. Finally, we found that native plant use diverged between landscapes, such that the probability of visiting a native flower increased over time in non-developed land uses but declined over time in developed ones, demonstrating that hummingbirds track the advancement of native floral phenology and use non-native, cultivated flowers as a human subsidy due to either availability or preferences.
4. Our preference and network models revealed that while hummingbird-plant network structure was similar across landscapes, the composition of important taxa shifted from native, wild species like *Monarda* and *Impatiens* to non-native, cultivated species like *Salvia*.
5. Using trait-based models of flower visitation, we found that hummingbirds preferred native, tubular, and red/orange flowers fitting the hummingbird pollination syndrome despite visiting >260 different plant species. Red and orange flowers were preferred across all seasons, suggesting color may be a reliable signal of nectar availability across species and contexts. Native and tubular flowers were strongly preferred during the breeding season, however, preferences relaxed during spring and fall migration.
6. These findings reveal the consistent preferences of Ruby-throated Hummingbirds for native, tubular, and red/orange flowers, and underscore how spatial and temporal factors reshape foraging behavior and trait preferences. Our results also highlight the value of community-collected data in characterizing plant-pollinator interactions across broad spatial and temporal scales.

## INTRODUCTION

Hummingbirds (family: Trochilidae) are essential pollinators for many plant species and are a well-studied model for the evolution of mutualisms between plants and pollinators (Ratto et al. 2018). Yet, even for common and well-studied species, information about plant-pollinator networks is limited to a few systems and typically focuses on natural ecosystems (Archer et al. 2014). Significant gaps exist in our understanding of plant-hummingbird interactions and how these networks vary across spatial, temporal, and land use gradients. Understanding how pollinators use resources across these gradients is essential for predicting how plant-pollinator networks may respond to habitat change. Thus, regional-scale studies that collect data across larger regions are needed to determine how hummingbirds and other pollinators interact with plants and other essential resources (Hazlehurst et al. 2021). Since some species of hummingbirds occupy wide distributions, are commonly observed by the public, and interact with both natural and anthropogenic resources, they represent an ideal group to explore these interactions with community-collected data.

Hummingbirds are among the most popular birds that capture public interest across communities (Schuetz and Johnson 2019; Andrade et al. 2022). Many species can also be frequently encountered around human settlements, making them easier to observe. Consequently, hummingbirds are some of the most frequently observed species on the iNaturalist platform (www.inaturalist.org), a community science website designed to collect large numbers of occurrence data points on living things. The growth and popularity of iNaturalist have enabled researchers to access presence-only occurrence data at larger spatial and temporal scales than ever before (Mason et al. 2025). Importantly, iNaturalist users often photograph organisms engaged in specific behaviors, and the platform allows annotation of interactive events, yield complementary information about how these organisms interact with their environment and with other species (Fonturbel et al. 2023). In the case of pollinators, observations can be leveraged to examine variations in resource use (Saldivar et al. 2022), such as visitation to both natural and cultivated floral resources and artificial feeders, making the data well-suited for studying plant-pollinator networks.

The Ruby-throated Hummingbird (*Archilochus colubris*, hereafter: ruby-throat) is the only native breeding hummingbird species of the Northeastern US. It is a Nearctic-Neotropical migrant and breeds throughout the eastern United States, and winters in Mexico and Central America (Weidensaul et al. 2020). Like other hummingbird species, ruby-throats feed on nectar from flowering plants for carbohydrates (Brice 1992) and also use pollen and insect resources for protein (Stiles 1995, Spence et al. 2022) to fuel their high metabolism. They can also use alternative resources, such as tree sap (Southwick & Southwick 1980) suggesting a generalist and flexible diet. They are found in a variety of habitat types and in developed and undeveloped landscapes, such as forests, forest edges, meadows, and gardens (Weidensaul et al. 2020). This habitat and resource generalism, combined with their willingness to use human-provided feeders, makes ruby-throats an ideal species for examining how resource use varies across natural and developed landscapes.

Previous studies of ruby-throat foraging ecology have documented preferences for large, tubular flowers and red and orange flowers (Binnie 1965, Bertin 1982), which are purported to have high concentrations of nectar relative to other flower colors (Rodríguez-Gironés and Santamaría 2004). Documented native flower preferences of ruby-throats in the far northwestern part of their range include trumpet creeper (*Campsis radicans*), coral honeysuckle (*Lonicera sempervirens*), scarlet bee balm (*Monarda didyma*), red columbine (*Aquilegia canadensis*), fire pink (*Silene virginica*), and red buckeye (*Aesculus pavia*) (Austin 1975, Bertin 1982). Ruby-throats are also reported to visit non-native flowers including Japanese honeysuckle (*Lonicera japonica*), butterfly bush (*Buddleia davidii*), and tropical sage (*Salvia coccinea*) (Binnie 1965). In a captive setting, ruby-throats selected flowers with longer corollas, higher nectar volume, and higher nectar concentration suggesting ruby-throats can accurately assess resource quality (Montgomerie 1984).

Despite how common ruby-throats are in the Eastern United States, assessments of their flower preferences and resource use are limited to just a few local and captive studies. None of these studies have evaluated ruby-throat flower preferences across a significant portion of their range or assessed how resource use varies over space and time (Hazlehurst et al. 2021). Moreover, these studies primarily occurred within natural ecosystems with wild, spontaneously occurring plants and have not considered how resource use might differ in developed land uses where many plants are cultivated (Maruyama et al. 2019). Lastly, although ruby-throats are known to use hummingbird feeders, it is unknown how feeder use might vary over space and time (Wolf et al., 2023). Understanding the relative importance of feeders versus native and non-native floral resources across different landscapes and seasons is critical for assessing whether anthropogenic subsidies are supporting or potentially disrupting natural plant-pollinator interactions.

In this study, we leveraged community-collected data from iNaturalist to conduct the first large-scale assessment of feeder and flower visitation patterns in ruby-throats across the Northeast (Figure 1). Specifically, we tested the hypothesis that ruby-throat resource use varies spatially, temporally and by land cover. To address this, we analyzed over 8 million occurrence records of hummingbirds and flowering plants, with over 2,100 annotated interactions between ruby-throats and flowers, as well as over 3,000 feeder visitation records. This dataset enables analyses at macroecological scales that would be challenging to amass via traditional field studies alone. We predicted that feeder use and flower use would vary spatially and temporally, such that feeder use would be more often observed in developed landscapes during spring and fall migration. We also predicted that plant use would be consistent across land uses, with more observations of visits to non-native, cultivated plants in developed land uses.

**Figure 1.**
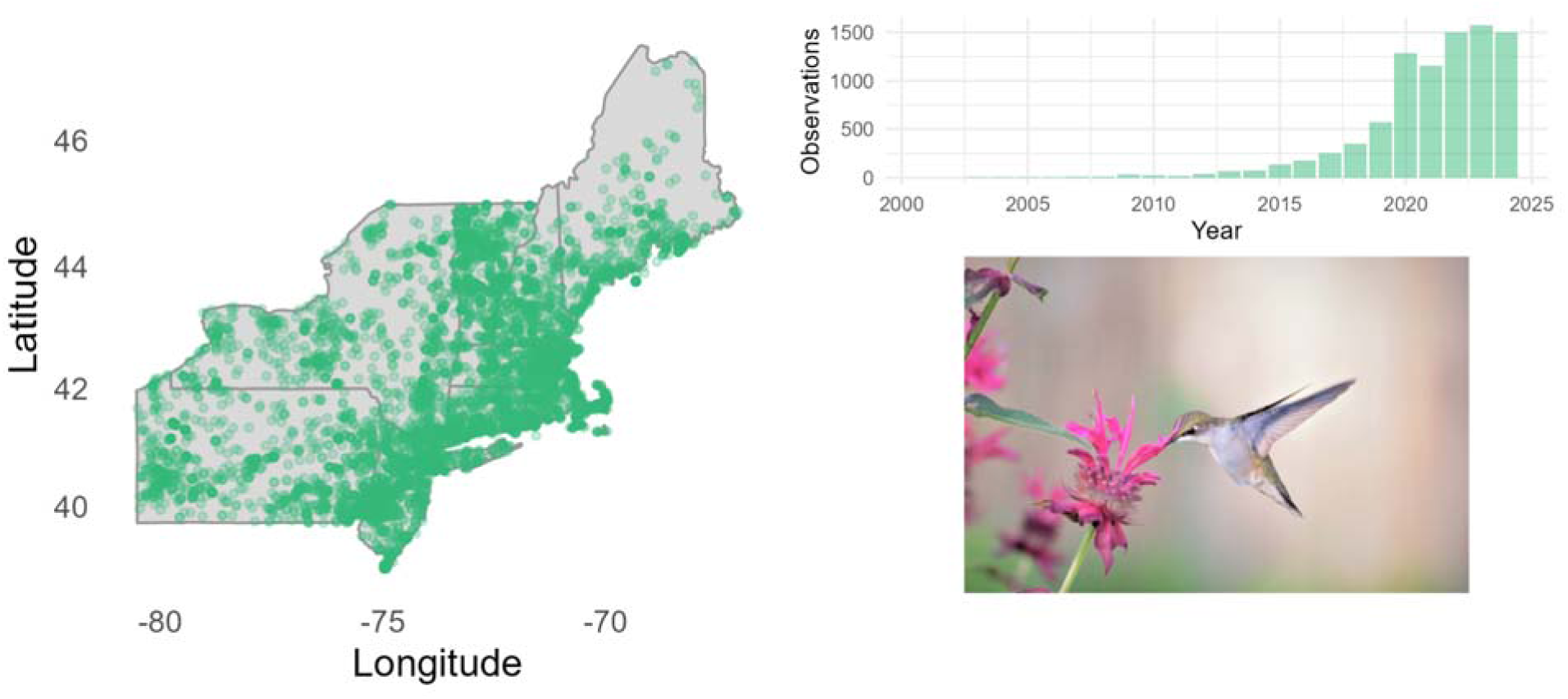
(A) Geographic distribution of >9k *iNaturalist* Ruby-throated Hummingbird observation across the study region. Each green point represents an individual observation. (B) Bar plot showing the increase in Ruby-throated Hummingbird observations over the study period (2000-2024), primarily from iNaturalist volunteer submissions (C). Example photo submission of a hummingbird visiting a flower (*Monarda didyma*).

Regarding flower preferences, we predicted that ruby-throats would have more observations visiting native plants relative to their availability. However, this preference would be weaker in developed areas and during the spring and fall migrations, when native flower abundance is more limited. Additionally, we predicted that ruby-throats would favor red and orange flowers, as well as tubular-shaped flowers, over other colors and shapes regardless of context. Here, we share our findings and offer insights into the resource use and flower species most preferred by these birds using community-collected data at regional scales.

## MATERIALS AND METHODS

### Ruby-throat observations

We reviewed 9,142 iNaturalist observations of ruby-throats from the Northeastern United States between 2023 and 2024. We defined the Northeast as the following states: Maine, New Hampshire, Vermont, Massachusetts, Connecticut, Rhode Island, New York, New Jersey, and Pennsylvania. We restricted analyses to observations with a known location in this region. We coded all observations as 1) flower visitation, 2) feeder visitation, or 3) no flower or feeder visitation. Flower visitations were defined as visual evidence of a hummingbird visiting or inserting its bill into a flower and were coded using the iNaturalist interaction field as ‘Interaction->visited flower of.’ Feeder visitations were defined as visual evidence of a hummingbird sitting on or feeding from a hummingbird feeder and were coded as ‘Visit to Bird Feeder.’ We considered observations of a hummingbird oriented toward a flower or feeder as evidence of visitation. All other observations were photographs of hummingbirds not engaging in foraging behavior, for example, perched, flying, preening, brooding young, or dead, among other examples.

We reviewed all observations of hummingbirds visiting flowers to identify plant species to the lowest taxonomic level, typically genus or species. The lead author verified plant identifications made by iNaturalist observers for accuracy. For plant taxa that are challenging to identify to species from photographs (e.g., *Solidago*), we identified them to genus or family. Observations of unknown plant taxa (in most cases due to poor picture quality) were recorded as visits to unidentified flowering plants. We consulted with botanists and online resources for assistance in identifying challenging plant species, typically cultivated varieties. Although some plants were identified by the observer at the cultivar or subspecies level, we did not include that information in our analyses.

We downloaded all observations directly from iNaturalist on November 30, 2024. Our final dataset included observations collected between 1989 and 2024 within our defined state boundaries. We initially sought to combine flower visitation records with those from the GloBi website (Poelen et al. 2014); however, 99.99% of flower-visit interactions were derived from our direct coding of iNaturalist observations rather than pre-existing GloBI records. Because some observers restrict data transfer from iNaturalist to other data portals, we downloaded all observations directly from iNaturalist through our community engagement project to maximize the observation records.

### Data Filtering

To reduce pseudoreplication, we first used a coarse-level filtering process to remove duplicate observations occurring at the same location, date, and visitation type (flower, feeder, or none). We also filtered observations with low spatial or temporal accuracy (e.g., those lacking precise month information). For temporal variation, we assigned each observation to one of three different seasons: spring migration (March, April and May), breeding Season (June, and July), and fall migration (August, September, and October). All observations collected in November, December, January, or February were considered vagrant individuals and discarded (n=21). For spatial variation, we used latitude to assess variation from south to north.

We classified whether observations occurred in developed or non-developed landscapes for each visitation. We used the 2023 National Land Cover Dataset (NLCD) provided at a 30-m pixel resolution. We extracted the land cover category for each observation point. We coded land cover as follows: developed-open space, developed-low intensity, developed-medium intensity, and developed-high intensity were coded as ‘developed,’ and all other land cover classifications were coded as ‘non-developed.’ We note that parks and rural residential areas with high tree canopy and very low housing density are typically classified as non-developed in this dataset, while urban, suburban, and exurban residential landscapes are typically classified as developed.

### Flower Availability

To assess flower preferences, we collected information on plant occurrences to account for availability. We extracted plant occurrence data from GBIF (GBIF.org, November 30, 2024). and supplemented these occurrences with ‘casual observations’ exported from iNaturalist. Casual observations are observations in iNaturalist that cannot reach research grade; in the case of plants, these are mainly cultivated species in gardens that were also strongly represented in our ruby-throat flower visitation records. Because we were interested in plant availability for wild and cultivated species, we combined these observations with the GBIF observations (which also includes research-grade iNaturalist observations). For all plants, we summed the number of observations of each genus as our measure of observation count. This accounted for the variation in how readily different genera are identified at the species level. Plant observations on community science platforms are a function of both abundance and observer behavior, therefore raw observations have very high variation and skew. Therefore, we used a scaled measure as our availability metric, scaled by the minimum and maximum values, so that availability values ranged between 0 and 1 with high values representing the most observed genera, and low values representing the least observed genera. We did not include genera that were never observed being visited by ruby-throats.

The relationship between abundance and observation frequency may be weaker for cultivated plant species for several plausible reasons: 1) observers may not submit cultivated plant observations that will never reach research grade, 2) observers with cultivated plant species in their gardens may not submit iNaturalist records, and 3) iNaturalist users may not submit records from developed landscapes with cultivated plants. In addition, within iNaturalist, research-grade observations are inconsistently applied to both wild and cultivated plants. To account for this discrepancy, we calculated two sampling metrics for each plant genus: the number of research-grade records (i.e., from GBIF) and the number of casual-grade records (from casual data submitted to iNaturalist). We used the plant observations to assign a cultivation status to each plant genus. For each genus, if a higher proportion of observations were casual grade, we classified the genus as primarily cultivated. If a higher proportion were research-grade, we classified the genus as primarily wild or naturalized.

### Flower Characteristics

For each hummingbird-flower interaction, we recorded attributes of each flower visited by ruby-throats. For each interaction, we recorded the plant family, genus, and species, the plant species origin (i.e., whether it was native to the Northeastern USA), flower shape, and its dominant flower color. For taxonomy, we used the plant taxonomy from GBIF. For plant origin, we used the BONAP database (Kartesz 2015) and classified plant species as native or non-native to the Northeast United States. To assign native and non-native to the genus level, we assessed the native status of each species or genus represented in our dataset.

For flower color, we assessed color for each individual photograph of a hummingbird-flower interaction (n=2,192). We used this approach, rather than classifying color at the species level, because many visited flower species had multiple color variants. To assess color, we used an AI approach through the OpenAI platform (OpenAI 2025). We developed a shell script to access the “gpt-4o-mini” vision model with the following prompt: “Can you identify the flower in this image and just tell me the color of the flower? Only report the color name.” We ran the script through a CSV file containing URLs linking to iNaturalist observation photos. This approach assigned color to 1,995 (91%) of our observations. For the remaining 197 (9%) observations, the model returned ‘unknown’ primarily due to poor image quality or instances where the observation had multiple photographs and the linked url did not contain a flower. For those observations, we manually assigned a color. For quality control assessments of the AI model, we manually annotated 195 observations and assessed the accuracy of the color assignment. 31/195 were incorrect (15%), with 6% error due to difficulties differentiating very similar colors (e.g., red versus orange), 3% of errors were due to unknowns (“I do not see a flower in this image”) and 6% provided the wrong color. Because we grouped similar colors (red and orange; purple and blue) and manually assigned color to all unknowns, we therefore report the accuracy of this model to be 94%.

We assigned flower shape using a similar approach but with the prompt: “text: Can you identify the flower in this image and just tell me the shape of the flower? Only report the shape name using these names: “radial”, tubular, funnel-shaped, bilabiate, radiate, grouped, umbellate, catkins, and spurred. We applied this model at the plant species level rather than to individual photographs because flower shape is not expressed variably within a species like color. The AI model assigned flower shape to 100% of plant species. Multiple authors reviewed all 283 taxa flower shape assignments for accuracy. For species where flower shape was described with two categories (“tubular, funnel-shaped”), we assigned the species the first flower shape.

### Statistical Analysis

#### Visitation Probability

We used a logistic regression approach to test the hypothesis that ruby-throat feeder and plant visitation vary with space, time, and land cover. We created three binary response variables: 1) feeder observation, 2) plant observation, and 3) native plant observation. We coded each variable so that 1 = true and 0 = false. Our temporal predictor was a continuous variable of calendar date between 60 (March 1st) and 304 (October 31st). Our spatial predictor was latitude, a continuous variable between 39° N and 45° N. Our land cover predictor was a categorical variable with two levels: developed and non-developed. We ran a generalized linear mixed model (GLMM) with year as a random intercept to account for annual variation in the observation process and a binomial distribution with a logit link. We ran a separate model for each response variable. We used the function “*glmer*” from the package lme4 (Bates et al. 2015) to run this model. We scaled date and latitude variables to the mean for model convergence.

#### Flower Species Preferences

To assess ruby-throat flower preferences, we modeled visitation counts for each plant genus as a function of plant availability, latitude, and land cover. We did not use season in this model, since our availability measures did not include a temporal component and plants can be observed regardless of whether they are flowering or not. We used iNaturalist observation frequency as a proxy for plant availability, assuming that observation frequency is broadly correlated with abundance. Because the distribution of plant observation frequency was strongly right skewed, we scaled the plant observation counts by the minimum and maximum to provide a relative metric of availability based on the analyzed dataset. We fit a negative binomial GLM with fixed effects of scaled plant observation frequency, latitude band (Mid-Atlantic/NYC [<40°N], southern New England [40–43°N], and northern New England [43+°N]), and land cover type (developed vs. natural). We defined preferred genera as those with large positive residuals, indicating visitation exceeding what would be expected given their availability and landscape context, and avoided genera as those with large negative residuals. We identified the most preferred genera within each latitude band and land cover type.

##### Flower Trait Selection

To determine whether ruby-throats selected for certain characteristics of flowers, we used a trait-based model. First, we assigned the following plant characteristics to each observation in the ruby-throat flower visitation dataset: flower shape (13 categorical levels), flower color (13 categorical levels), native status (2 levels; native and non-native; unknown was excluded), genus (character variables), cultivated status (2 levels), and relative abundance (scaled between 0 and 1 using min-max scaling). We then summarized ruby-throat visitation counts so that our response was the total interactions for each combination of flower shape, flower color, and native status across temporal (Spring migration, Breeding, Fall Migration) and landscape (developed and natural) contexts. We did not include region to reduce overparameterization and because we did not predict a mechanism by which trait selection would vary across the region.

We next classified each trait combination as fitting the hummingbird pollination syndrome using binary variables. Flowers with tubular or bilabiate shapes were classfied as tubular (1 vs. 0). Flowers that were red or orange, were classified as red/orange (1 vs. 0), and flower genera that were native, were classified as native (1 vs. 0). We fit a negative binomial generalized linear model with visitation count as the response, and fixed effects of scaled plant availability, tubular, red/orange, native, season, and land cover, as well as two-way and three-way interactions between these variables (excluding scaled abundance, which was included as a covariate). Non-significant interactions were dropped from the final model. We used a negative binomial distribution to account for overdispersion in the response variable.

#### Hummingbird-flower Network Analysis

To assess whether ruby-throat interaction networks differed between developed and non-developed landscapes, we constructed two weighted bipartite networks where one set of nodes represented plant genera, and the other set represented ruby-throats in different spatiotemporal contexts (season × latitude band combinations). Because we only had one bird species in this dataset, we treated ruby-throat populations in different spatiotemporal contexts as functionally distinct network nodes (e.g., “Northern New England – Fall”), allowing us to examine resource use across landscape contexts. This approach follows the framework of population-level interaction networks, which recognizes that intraspecific variation in resource use across space and time can be analyzed using network metrics typically applied to multispecies communities (Poisot et al. 2015). We created separate networks for developed and non-developed land cover types, each containing up to nine spatiotemporal context nodes (3 seasons × 3 latitude bands). We weighted links by the number of observed visitation interactions between each spatiotemporal context and each plant genus.

We calculated network-level metrics including connectance (the proportion of realized interactions), nestedness (NODF), modularity, and network-level specialization (H2’). We also calculated genus-level metrics including normalized degree (accounting for differences in network size) and genus strength (the sum of dependencies of all partners). We compared network metrics between developed and non-developed landscapes using permutation tests (1000 iterations). To assess whether observed network structures differed from random expectations, we used null models that randomized interactions while maintaining network dimensions. We visualized networks using bipartite plots ordered by interaction frequency. All network analyses were conducted using the bipartite package (Dormann et al. 2008) in R.

## RESULTS

### Observation Summary

Our final dataset included 9,125 coded observations of ruby-throats extracted from *iNaturalist*, of which more than half of the observations were feeding events. Visits to feeders accounted for 2,797 of all observations (31%), with 42%, 35%, and 24% occurring in spring migration, breeding season, and fall migration, respectively. 2,125 (22%) of observations were to flowers, with 8%, 30%, and 62% occurring in spring migration, breeding season, and fall migration respectively. The remaining 47% of observations were of non-feeding events.

### Resource Use

#### Feeders

We found that the probability of ruby-throats feeder visitation probability varied by season, land cover, and the interaction between season and land cover (Table 1, Figure 2). The probability of a feeder visitation was highest during spring migration and lowest during fall migration. The probability of visiting a feeder was consistently higher in developed landscapes than in non-developed landscapes, and this difference was greatest during spring migration and weakest during fall migration.

**Figure 2.**
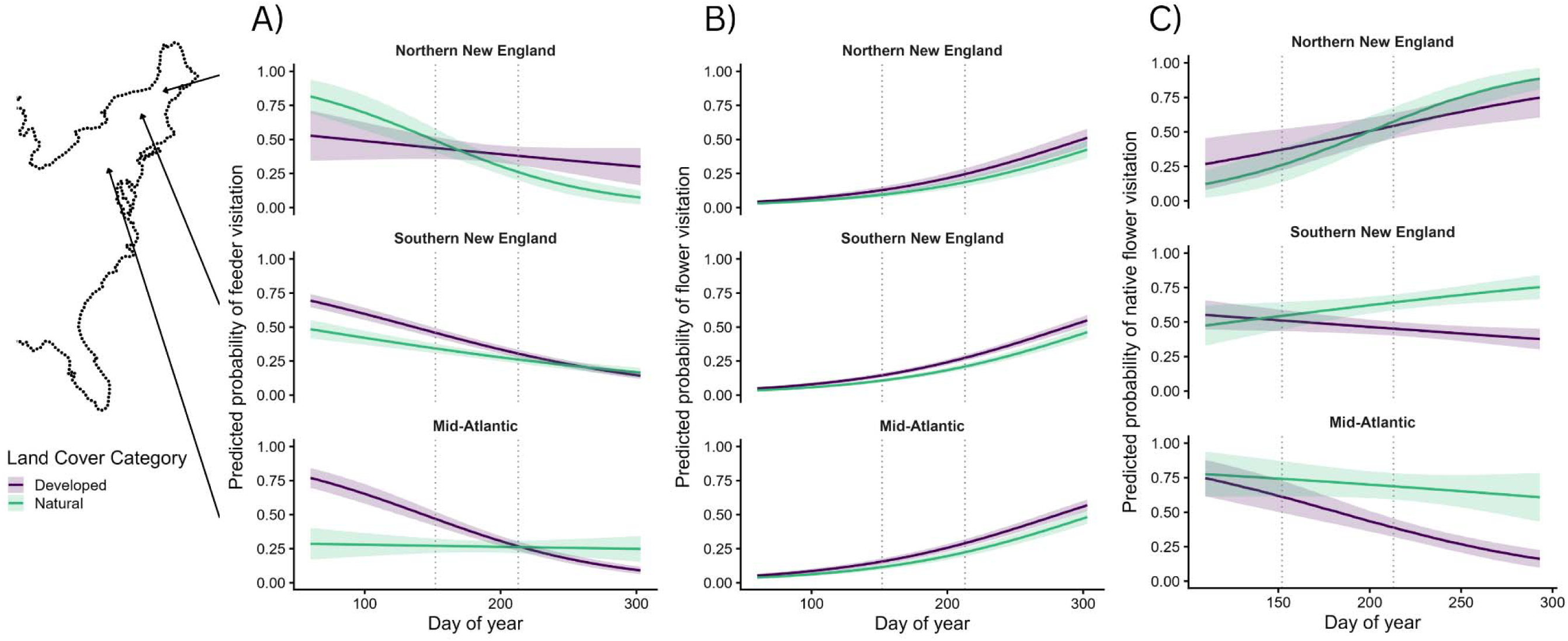
Probability of (a) feeder visitation, (b) flower visitation, and (c) native flower visitation by Ruby-throated Hummingbirds across calendar date, land use (developed vs. undeveloped), and latitude. The logistic regression model included latitude as a continuous variable, but predictions are shown for three illustrative levels representing the regional bands: the Mid-Atlantic (39), Southern New England (42), and Northern New England (45). Colors correspond to land use categories, where developed land uses are shown in purple, and natural land uses in green. Lines represent model predictions, shaded areas indicate 95% confidence intervals. Vertical dotted lines represent the transition from spring (left), summer (middle), and fall (right) seasons.

**Table 1.** Results from logistic regression models examining the probability of feeder, flower, or native flower visitation by Ruby-throated Hummingbirds. Predictor variables include calendar date, latitude (scaled), land use (developed vs. undeveloped), and their interactions. Coefficients (β), standard errors (SE), z-values, and p-values are reported for each predictor.

| Response | Predictors | $\beta \pm SE$ | *z* | *P* |
| --- | --- | --- | --- | --- |
| *Feeder Visitation* | Intercept | -0.69 ± 0.07 | -10.29 | <0.0001 |
| | Date | -0.49 ± 0.03 | -15.44 | <0.0001 |
| | Latitude | 0.07 ± 0.03 | 2.21 | 0.03 |
| | Land Use (Undeveloped) | -0.27 ± 0.05 | -5.19 | <0.001 |
| | Latitude : Land Use (Undeveloped) | -0.04 ± 0.05 | -0.70 | 0.48 |
| | Latitude : Date | 0.09 ± 0.03 | 2.79 | 0.005 |
| | Date : Land Use (Undeveloped) | 0.20 ± 0.05 | 3.81 | 0.0001 |
| | Date: Latitude : Land Use | -0.22 ± 0.05 | -4.32 | <0.0001 |
| *Flower Visitation* | Intercept | -1.13 ± 0.05 | -20.74 | <0.0001 |
| | Date | 0.59 ± 0.03 | 20.28 | <0.0001 |
| | Latitude | -0.04 ± 0.03 | -1.60 | 0.11 |
| | Land Use – Undeveloped | -0.34 ± 0.06 | -6.15 | <0.0001 |
| *Native Flower Visitation* | Intercept | -0.18 ± 0.11 | -1.69 | 0.09 |
| | Date | -0.12 ± 0.06 | -1.98 | 0.05 |
| | Latitude | 0.17 ± 0.06 | 2.87 | 0.004 |
| | Land Use (Undeveloped) | 0.82 ± 0.11 | 7.54 | <0.0001 |
| | Latitude : Land Use (Undeveloped) | -0.22 ± 0.10 | -2.15 | 0.03 |
| | Latitude : Date | 0.22 ± 0.05 | 4.19 | 0.001 |
| | Date : Land Use (Undeveloped) | 0.37 ± 0.11 | 3.29 | 0.002 |

#### Flower visitation

The probability of visiting a flower was related to season, latitude, land cover, and the interactions between these three predictors (Table 1, Figure 2). During spring migration, the probability of ruby-throats visiting flowers was very low at all latitudes and land cover types. During the breeding season, the probability of visiting a flower was higher in developed landscapes at low latitudes and declined as latitude increased. During fall migration, there was a strong interaction between land cover and latitude, such that flower visitation was higher in developed landscapes at low latitudes but higher in non-developed land uses at higher latitudes.

#### Native Plants

When comparing only visits to plants, we found a relationship between native plant visitation and our three predictors (Table 1, Figure 2). During spring migration, native plant visitation declined with latitude, and this negative relationship was strongest in developed landscapes, where the probability of native plant visitation approaches zero above 43 degrees latitude in the spring (approximately southern Maine). During the breeding season, the probability of visiting a native plant decreased with latitude in both developed and non-developed land uses. That decrease was steeper in non-developed landscapes shifting from 0.85 at the lowest latitudes to nearly 0.12. During fall migration, the probability of visiting a native flower was consistently higher in non-developed land uses but declined with latitude. Whereas native visitation was low in developed land uses consistently across entire latitudinal gradient.

### Flower and trait preferences

Our dataset included 2,125 unique interactions of ruby-throats with 264 flowering plant species representing 158 genera. 99.43% of photos could be identified to at least genus, 82.21% to species, and 2.68% (n=57) records were identified as hybrid taxa. 0.56% (n=12) records were identified at a coarser taxonomy (family or higher) or remained unidentified. These records identified were excluded from further analyses.

Ruby-throat visitation patterns to plant species differed between developed and non-developed landscapes. In developed landscapes, hummingbirds were observed visiting 218 different plant taxa with 1,384 interaction observations, while in non-developed sites, they visited 153 plant taxa across 603 interaction observations. 111 plant taxa were only observed in developed landscapes, while 48 were only observed in natural landscapes.

#### Flower Preferences

Across the Northeast US, the most visited plant species in spring migration was *Rhododendron* sp; during the breeding season, it was *Monarda didyma*; and during fall migration, it was *Impatiens capensis*. In natural landscapes, interactions were dominated by *Monarda didyma* (16%), *Impatiens capensis* (15%), and *Lobelia cardinalis* (13%). In developed landscapes, interactions were dominated by *Salvia guaranitica* (11%), *Lobelia cardinalis* (10%), and *Monarda didyma* (10%).

According to our flower preference model, visitation frequency to a plant genus was significantly predicted by availability (β =19.15 ± 6.35 SE), region (Northern New England: β =-8.05 ± 2.64; Southern New England : β =-6.60 ± 2.55), land cover (Natural: β = -8.31 ± 2.62) and the interaction between region and land cover (Northern New England X Natural: β =8.69 ± 3.77; Southern New England X Natural : β = 7.38 ± 3.63). Genera with the largest positive residuals, indicating visitation exceeding expectations given availability, were predominantly native species, along with a few non-native genera (Table 2, Figure 3). Across all three regions and both land cover types, *Monarda* (mean residual 39.83 ± 17.5 SD), *Lobelia* (28.5 ± 38.7), and *Impatiens* (21.6 ± 13.5) were consistently preferred. *Salvia* was an outlier in the dataset, with high average preference but extreme variation across contexts (53.9 ± 108), likely driven by high visitation in developed landscapes but absence in natural ones.

**Figure 3.**
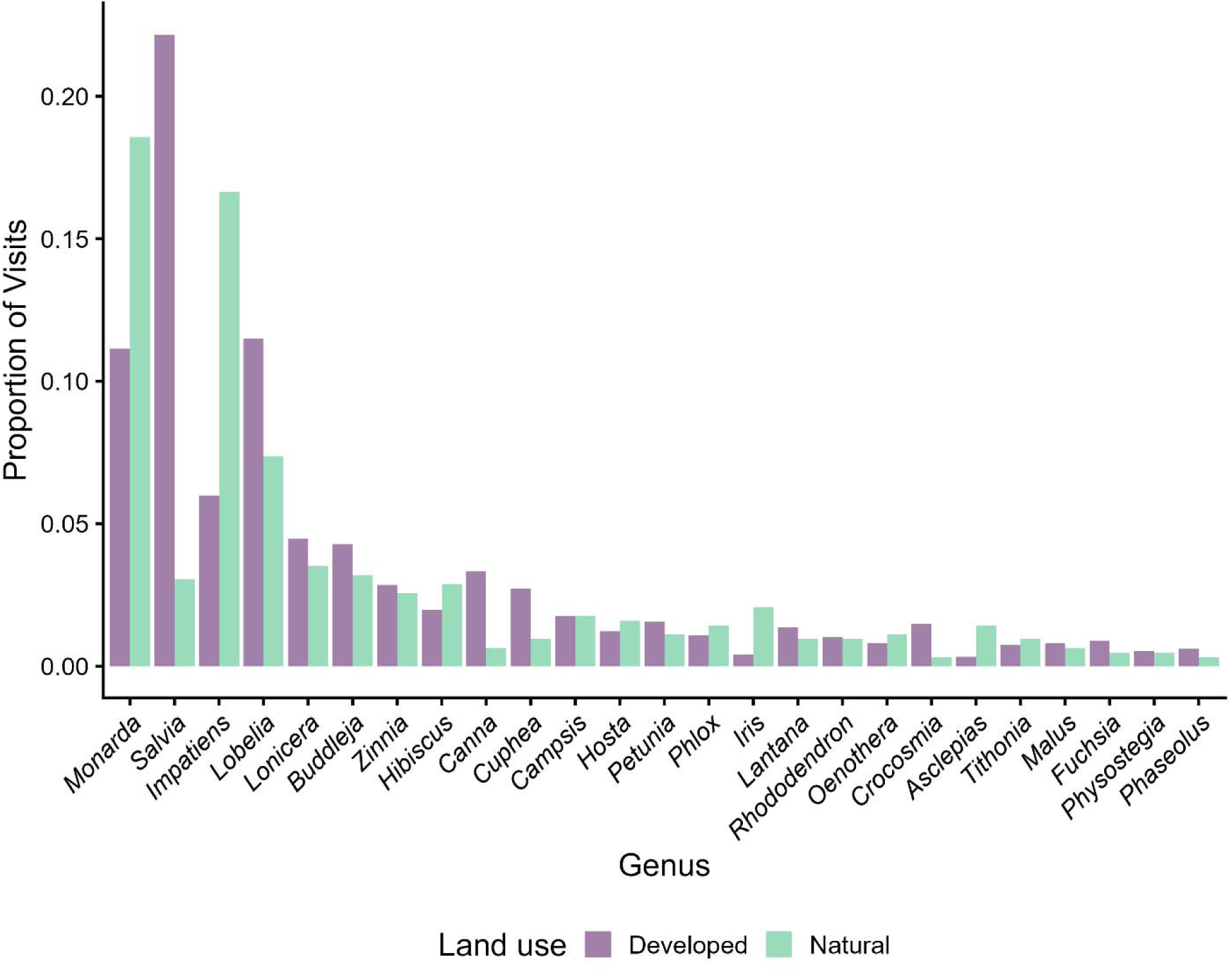
Proportion of total observations by plant taxon and land cover for the most visited genera in the dataset. Bars represent the raw counts of observations across the entire region. Only genera with >10 total observations are included.

**Table 2.**
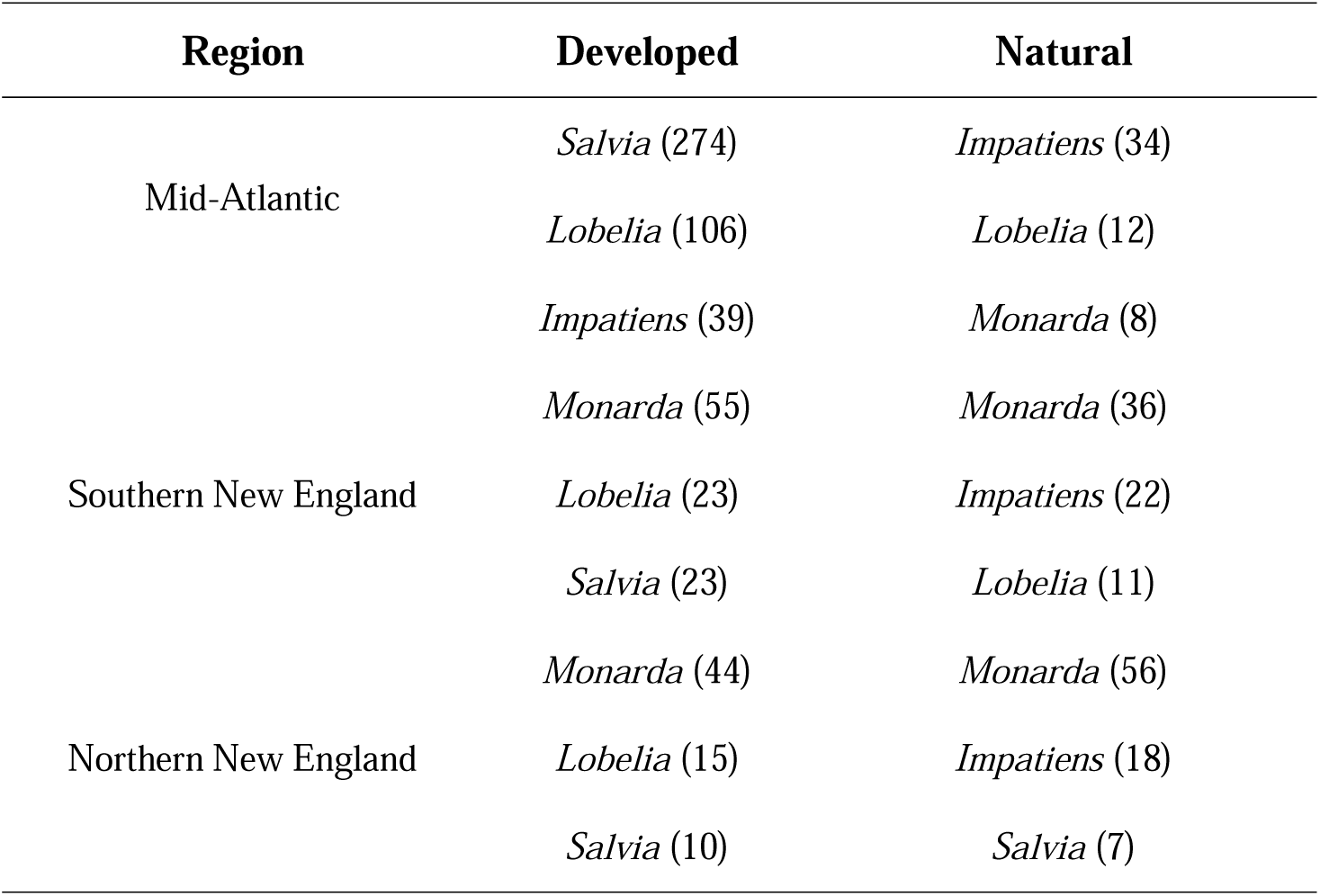
Table of flower genera with the highest preference values for each region and each land cover type (developed or natural). Numbers in parentheses represent the residual value from the flower preference model.

### Network Comparison

Using our population-specific network comparison, we found that network structure was not significantly different (P>0.1) between developed and natural landscapes, but the identities of important nodes varied (Table 3, Figure 4). Both networks exhibited higher modularity and network specialization than expected by random chance (P<0.001). Although network structure was similar, evenness and relative importance differed. The natural network had higher species strength, suggested less evenly distributed interactions across the network, compared to networks in developed landscapes (Table 3). The top plant genera were similar across both networks, however, more cultivated, non-native species represented in the developed network, and interactions were distributed more evenly across taxa (Table 3).

**Figure 4.**
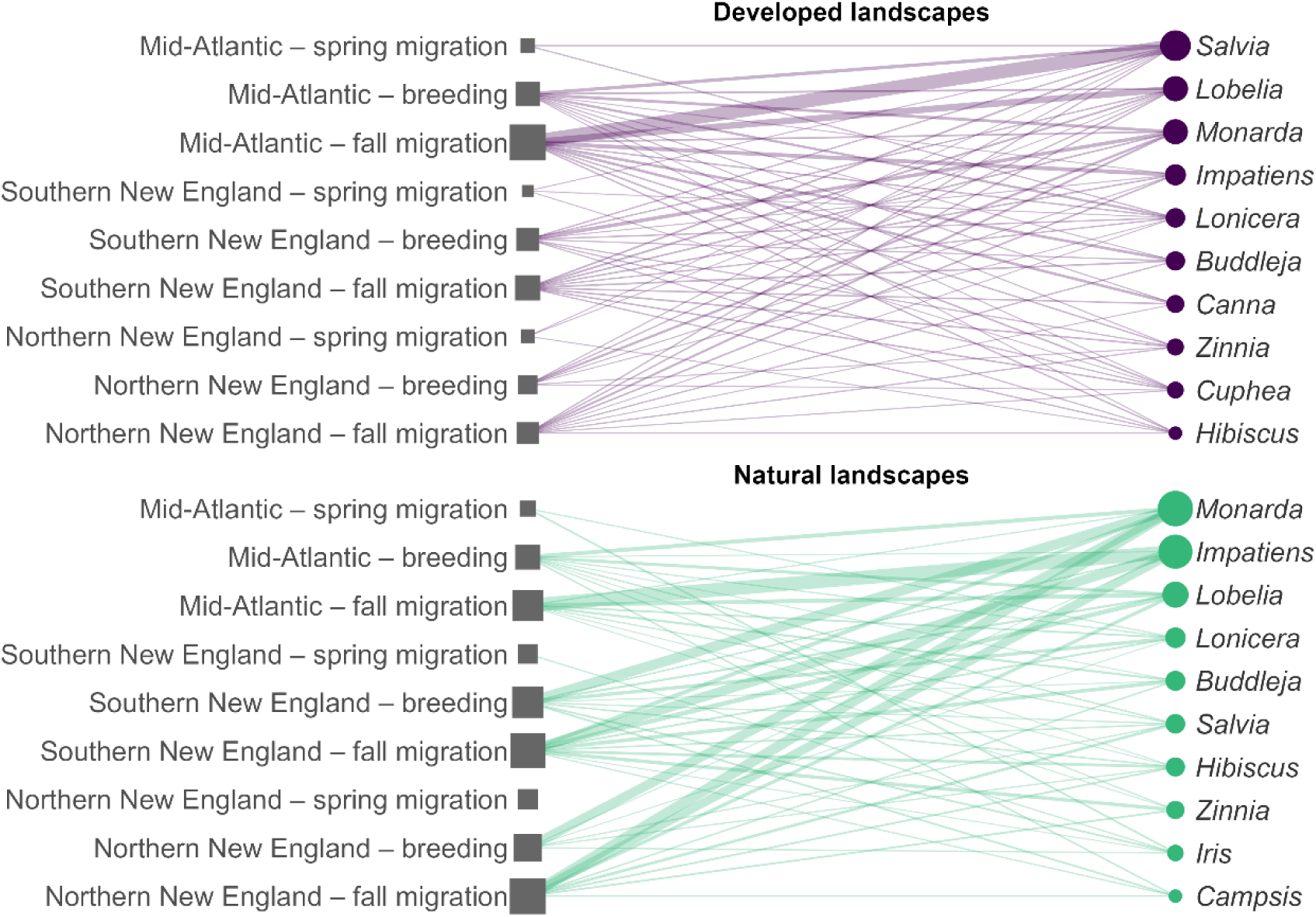
Bipartite network graph of interactions between ruby-throat population nodes (Region – Season) and plant genera in developed (purple) and natural (green) landscapes. Node size is proportional to visitation. For clarity, only the top 10 plant genera for each network are shown.

**Table 3.** Genus-level metrics of the top 10 plant genera in natural and developed networks.

| Network | Genus | Normalized Degree | Species Strength | Origin | Type |
| --- | --- | --- | --- | --- | --- |
| Natural | *Monarda* | 0.67 | 1.32 | Native | Wild |
|  | *Impatiens* | 0.67 | 0.96 | Native | Wild |
|  | *Lobelia* | 0.55 | 0.50 | Native | Wild |
|  | *Iris* | 0.55 | 0.43 | Both | Wild |
|  | *Lonicera* | 0.67 | 0.36 | Both | Wild |
|  | *Elaeagnus* | 0.33 | 0.29 | Non-native | Wild |
|  | *Aesculus* | 0.22 | 0.22 | Native | Wild |
|  | *Vaccinium* | 0.33 | 0.21 | Native | Wild |
|  | *Rhododendron* | 0.55 | 0.21 | Unknown | Wild |
|  | *Salvia* | 0.67 | 0.20 | Non-native | Cultivated |
| Developed | *Salvia* | 1.00 | 1.17 | Non-native | Cultivated |
|  | *Monarda* | 0.67 | 1.09 | Native | Wild |
|  | *Lobelia* | 0.77 | 0.62 | Native | Wild |
|  | *Lonicera* | 0.88 | 0.50 | Both | Wild |
|  | *Rhododendron* | 0.55 | 0.41 | Unknown | Cultivated |
|  | *Malus* | 0.44 | 0.34 | Unknown | Cultivated |
|  | *Impatiens* | 0.67 | 0.31 | Native | Wild |
|  | *Buddleja* | 0.55 | 0.22 | Non-native | Cultivated |
|  | *Zinnia* | 0.67 | 0.21 | Non-native | Cultivated |
|  | *Cuphea* | 0.77 | 0.20 | Non-native | Cultivated |

### Flower Trait Preferences

After controlling for plant availability, ruby-throats showed strong preferences for tubular flowers (β = 0.62 ± 0.16 SE, *P* < 0.001), red or orange flowers (β = 0.62 ± 0.10, *P* < 0.001), and native species (β = 0.74 ± 0.16, *P* < 0.001) during the breeding season (Table 4, Figure 5). Trait preferences did not differ between developed and natural landscapes (β = 0.11, SE = 0.09, p = 0.21), indicating that hummingbirds maintain consistent floral trait preferences regardless of context. However, trait selectivity varied seasonally such that preference for native species declined significantly during both fall (β = -0.63 ± 0.20, p = 0.002) and spring migration (β = -0.85 ± 0.30, *P* = 0.005). Preference for tubular flowers persisted during fall migration but was lower during spring migration (β = -0.83 ± 0.34, *P* = 0.01), when floral sources are limited. Red and orange flower preference remained consistent across all seasons. Similar to the flower visitation results, visitation was highest during fall migration (β = 0.77 ± 0.18, *P* < 0.001), likely reflecting increased foraging when floral availability is high.

**Figure 5.**
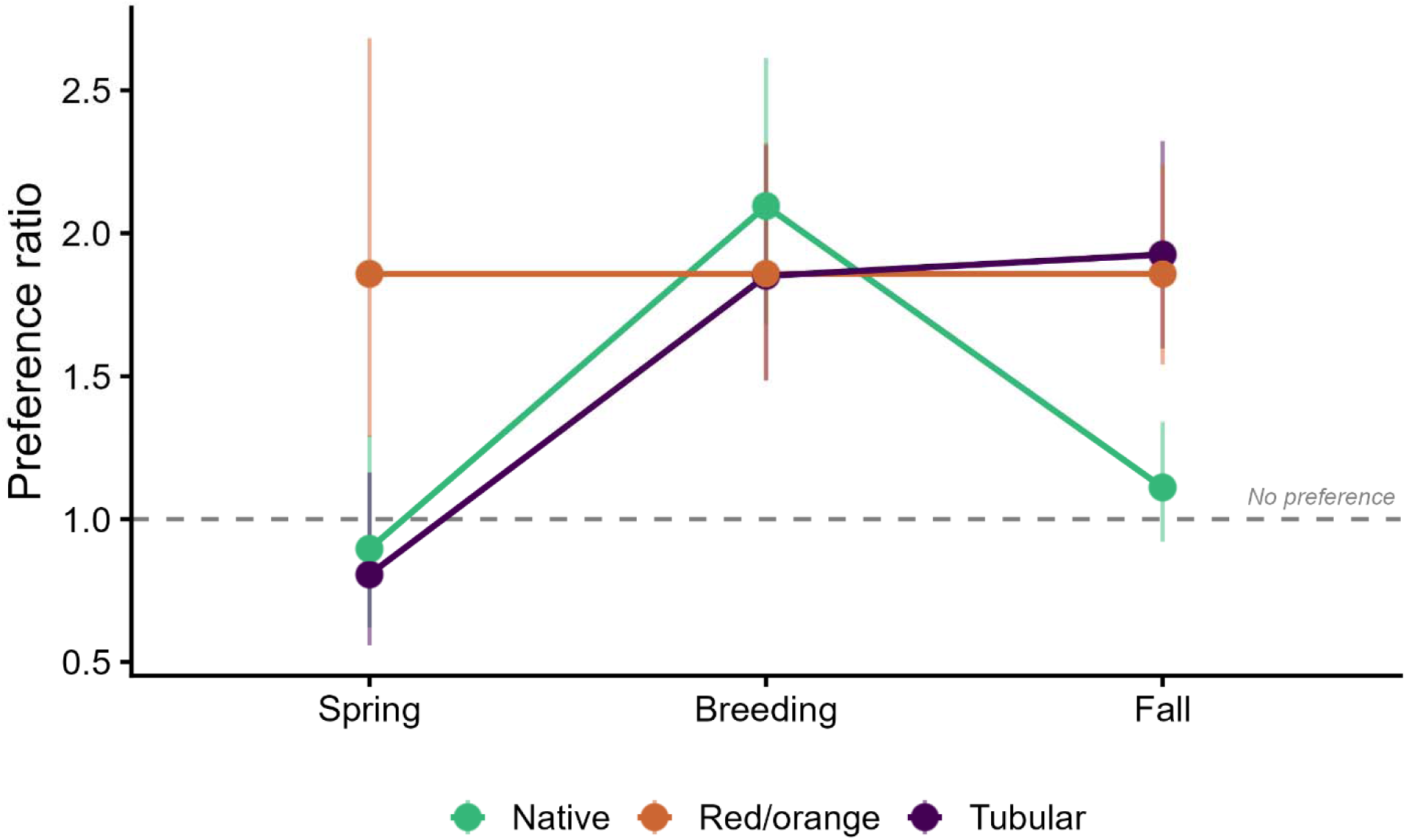
Seasonal variation in ruby-throat floral trait preferences. Points show the preference ratio (predicted visitation to flowers with a trait divided by predicted visitation to flowers without that trait), derived from a negative binomial GLM controlling for plant availability. Values above the dashed line (ratio = 1) indicate preference; values at or below indicate no preference. Error bars show 95% confidence intervals.

**Table 4.** Results of negative binomial GLM predicting ruby-throat flower visitation counts as a function of plant availability, floral traits, and season.

| Predictor | Estimate | 95%<br>CI | *z* | *P* |
| --- | --- | --- | --- | --- |
| Intercept | 0.16 ± 0.15 | -0.14, 0.46 | 1.02 | 0.306 |
| Scaled abundance | 1.13 ± 0.25 | 0.47, 1.81 | 4.61 | < 0.001 |
| Tubular | 0.62 ± 0.16 | 0.29, 0.94 | 3.81 | < 0.001 |
| Red/orange | 0.62 ± 0.10 | 0.41, 0.83 | 6.01 | < 0.001 |
| Native | 0.74 ± 0.16 | 0.43, 1.06 | 4.74 | < 0.001 |
| Natural land cover | 0.11 ± 0.09 | -0.07, 0.29 | 1.26 | 0.206 |
| Fall migration | 0.77 ± 0.18 | 0.43, 1.11 | 4.32 | < 0.001 |
| Spring migration | 0.44 ± 0.30 | -0.13, 1.05 | 1.49 | 0.136 |
| Tubular × Fall migration | 0.04 ± 0.20 | -0.36, 0.45 | 0.20 | 0.845 |
| Tubular × Spring migration | -0.83 ± 0.34 | -1.51, -0.18 | -2.47 | 0.013 |
| Native × Fall migration | -0.63 ± 0.20 | -1.03, -0.24 | -3.17 | 0.002 |
| Native × Spring migration | -0.85 ± 0.30 | -1.44, -0.25 | -2.79 | 0.005 |

## DISCUSSION

### Summary

Our results yield new insight into how ruby-throat resource use varies over space, time, and land use, demonstrating the complexity of these relationships. Feeder visitation was highest during spring migration, particularly in developed land uses, and declined steadily over time, reaching its lowest levels during fall migration. In contrast, flower visitation increased across the study region, with more visits in developed land uses and the highest visits during fall migration. Regarding native versus non-native plant use, we observed high and increasing visitation to native plants in undeveloped land uses across the region and in developed land uses in northern New England. However, native plant visitation sharply declined over time in developed land uses of more southerly regions (Southern New England, Mid-Atlantic). For flower visitation, we found that although ruby-throats visited a large diversity of different flowers, they exhibited high preferences for certain genera regardless of context. Similarly, network metrics were similar across landscapes. Lastly, while hummingbirds consistently preferred red and orange flowers, their preference for tubular flowers and native plants was seasonally dependent. Below, we explore potential mechanisms driving these patterns.

### Temporal

Our results provide new insights into the temporal shifts in resource use by Ruby-throated Hummingbirds. While it is well established that ruby-throats, like many hummingbirds, frequently visit feeders (Alexandre, et al. 2025), our findings indicate that feeder use is most common during spring migration, when the birds travel north to their breeding grounds and declines steadily over time. Controlling for time, it is also more common in the northern part of their range where flower phenology accumulates more slowly. In contrast, flower visitation increased as the season progressed across all latitudes. These patterns may reflect a tracking of floral phenology: hummingbirds rely on human-provided feeders when flowers are scarce early in the season, gradually shifting to floral resources as more species bloom. Feeders could serve as critical resource buffers during early migration or periods of harsh weather, such as unseasonably cold conditions in the spring. Additionally, earlier migration timing driven by climate change may increase reliance on feeders when floral resources are unavailable or facilitate range expansion into previously unsuitable habitats (Grieg et al. 2017, Alexandre, et al. 2025).

Alternatively, shifts in feeder use might reflect changing energetic needs. During spring migration, hummingbirds prioritize fat and sugar to fuel their long-distance flights. In contrast, floral resources and insect predation might better meet protein needs later in the season during breeding (Dick et al., 2020). However, the observed low feeder use during fall migration suggests that resource availability, rather than energetic requirements, may primarily drive these patterns. When floral availability is limited and competition the highest, hummingbirds may be more likely to take advantage of feeder subsidies (Zenzel and Moore 2019). Whichever the mechanism, feeder use appears to be a primary resource of spring migration in this species when floral availability is low.

### Landscape

Our findings show that land use is a strong predictor of resource use by Ruby-throated Hummingbirds. Both feeder and flower visitation were higher in developed land uses, suggesting these areas serve as important hubs of resource availability. Anthropogenic subsidies, such as feeders, refuse, and other human-provided foods, are more abundant in developed areas (Lerman et al. 2020). Additionally, residential landscapes and home gardens often feature a high diversity and abundance of floral resources cultivated primarily for their aesthetic appeal (Lerman et al. 2023). Some studies suggest that residential gardens may support greater plant diversity and abundance than nearby natural areas, due to cultivation of both native and non-native species (Padulles-Cubino et al. 2020). These developed landscapes may thus provide critical resources for hummingbirds, particularly during energetically demanding periods like spring migration.

However, the benefits of developed landscapes are accompanied by potential risks. These habitats may expose hummingbirds to anthropogenic threats, including outdoor cats, window collisions, and pesticide exposure (Loss and Marra, 2015). While floral resources are typically abundant in developed areas, it remains unclear whether these flowers meet the nutritional needs of hummingbirds. Nectar and pollen quality can vary significantly among plant species, and cultivated gardens often feature non-native species, hybrids, and cultivars (Padulles-Cubino et al. 2020), which may, or may not, offer reduced nectar or pollen availability, regeneration rates, or nutritional quality (Staab et al. 2020, Dorian et al. 2026). Importantly, nectar alone does not meet all of a hummingbird’s dietary requirements. Insects are a critical food source, particularly during the breeding season, providing essential protein and nutrients (Carroll et al., 2023). However, insect abundance tends to be lower in urban and developed landscapes (Lerman et al. 2021) and on non-native plants (Tallamy et al. 2021). This raises the possibility that abundant floral resources in developed areas may act as a “dishonest signal” of habitat quality, attracting hummingbirds to breed in environments with low prey availability or heightened threats. Future research should explore the demographic responses of hummingbirds in anthropogenic habitats, assessing resource availability, long-term survival, and reproductive success.

### Native and Non-native Plants

Complementing studies of other hummingbird species, our results reveal that ruby-throats use both native and non-native plants (Hazlehurst et al. 2021), however our study finds that native plants are generally preferred relative to availability. We also build on this by finding that this relationship diverges spatially and temporally. During spring migration, native plant use was highest in southern New England and lowest in northern New England, likely reflecting differences in spring phenology and the range limits of preferred plants. For example, *Monarda didyma* (Scarlet Bee Balm) and *Campsis radicans* (Trumpet Vine) are native to Mid-atlantic United States but do not regularly occur in the wild in New England.

In non-developed areas and in northern New England, native plant use increased or remained steady over time. However, in more southerly developed areas, the probability of using a native plant declined. This pattern could reflect differences in landscaping styles across latitudes. Native plants and spontaneous vegetation may be more common in the north, while non-native plants and cultivated vegetation dominate southern landscapes. Additionally, some highly preferred non-native plants, such as *Salvia* species, may have range limits in the northern US. Many non-native plants are selected for their floral abundance and attractiveness to pollinators, which could explain their high visitation rates in southern developed areas. However, despite these shifts, preference for natives was high during the breeding season in all contexts (Table 3, Table 3, Figure 5) and only declined during spring and fall migration when availability of highly preferred native species (*Monarda, Impatiens, Lobelia*) may be limited. Even during these periods, non-native plants were not strongly preferred relative to availability, despite frequent visitation. These findings suggest that resource availability and plant quality vary substantially across landscapes and regions.

### Flower Preferences

Our analysis of flower preferences aligns with established literature, showing that hummingbirds consistently favor tubular and red/orange flowers, a relationship shaped by coevolution of hummingbirds with plant species at a macroevolutionary scale. Several tubular and red/orange flowering species reported to be preferred by hummingbirds, including native Common Jewelweed (*Impatiens capensis*), Cardinal Flower (*Lobelia cardinalis*), and Scarlet Bee Balm (*Monarda didyma*), emerged as highly visited in our study (Table 2). Bilabiate *Salvia* species dominated visitation among non-native plants, underscoring their attractiveness to hummingbirds. Hummingbirds are thought to track the phenology of *Impatiens capensis* across its range (Bertin 1982), a pattern observed here in both developed and non-developed land uses. However, although tubular flowers are generally favored, they are not exclusively used, particularly during spring migration when availability is most limited. This generalism toward flowers that vary in shape that do not typically represent the ‘hummingbird syndrome’ suggest strong generalized mutualisms rather than highly specialized relationships like those found in other hummingbird clades (Abrahamczyk and Renner 2015, Rodríguez-Flores et al. 2019).

Interestingly, while preferences for tubular flowers diverged, color preferences was consistent between seasons and landscape context. Hummingbirds exhibited strong preferences for red and orange flowers, likely reflecting their superior nectar content and evolutionary resource availability signaling (Montgomerie 1984). Despite high visitation of blue and purple non-native flowers (particularly Salvia, a strong oulier in our dataset) when controlling for availability, red and orange flowers were still highly preferred almost 2:1. This consistent color preference may result from an honest signaling that remains despite artificial selection in cultivated plants. Alternatively, ruby-throats may select for red and orange flowers due to evolutionary adaptation even when higher nectar flowers of alternative colors are available. In developed landscapes, the availability of high-quality red and orange flowers may be reduced due to human preferences in landscaping for certain color palettes. While the evolutionary tie between flower color and ruby-throats appears robust, the weaker relationship between flower shape and visitation in non-native plants suggests that nectar availability and quality may play a more central role in hummingbird resource selection than evolutionary links to flower shape (Fenster et al. 2019, Lunau et al. 2011). Future research should explore how regional plant communities and human landscaping choices influence these relationships.

One caveat to the use of community-collected data is that our measures of cultivated flower availability may be conservative due to limitations in data availability. Here, we used occurrence data from GBIF coupled with research-grade and casual iNaturalist observations, which, to our knowledge, is the only available plant data of cultivated species at this scale. Our measure of preference (use relative to availability) may overestimate preferences for cultivated flowers due in part to the lack of information on their actual availability in the landscape. For example, Marigolds (*Tagetes erecta*), one of the most popular plants sold in nurseries, were represented by only 323 observations in our dataset, likely underrepresenting their actual occurrence in the landscape as an available foraging resource for hummingbirds. Thus, the high preferences for some non-native, cultivated plant species may also be an artifact of the low availability of occurrence data for these species. Our research highlights a need for efforts to systematically collect data on cultivated plant diversity in addition to spontaneous plant diversity at broader spatial scales and studies that assess macro-scale plant-pollinator data for relative comparisons between plants found in the wild and in cultivation.

## Conclusion

Our study provides important insights into the temporal, spatial, and land use patterns of resource use by Ruby-throated Hummingbirds. By highlighting the differences between feeder use, floral use, and floral preferences, we demonstrate how human-modified environments can both support and potentially hinder hummingbird populations. Our results also demonstrate how community-collected data can be used to elucidate ecological patterns at larger spatial and temporal scales (Marin-Gomez et al. 2022) and demonstrate the tremendous volume of observations that can be collected in relatively short periods of time. Indeed, the observations included here vastly increase the number of known ruby-throat-associated plants if relying on peer-reviewed literature or databases alone particularly for cultivated species, as has been shown in tropical systems (Marin-Gomez et al. 2022).

The patterns observed in this study highlight the need for future research in the individual and population-level responses of hummingbirds to land use change, anthropogenic threats, and shifting resource use. Moreover, although it is apparent that hummingbirds use non-native plants, they highly prefer native species. Further exploration into ruby-throat performance, resource quality, diet, and energetic needs is necessary to determine how and to what extent cultivated habitat in developed landscapes foster an important part of a conservation strategy for hummingbirds and other pollinators, or pose a population sink.

## ACKNOWLEDGEMENTS

This work was made possible by the millions of observations provided by community scientists that contribute to iNaturalist and other crowd-sourced projects. We specifically recognize the efforts of members of the project “*Pollinator Interactions on Plants*” Project who contributed to or annotated most of the observations used for this paper. We also acknowledge the contributions of Amelia Dubois whose efforts helped with the feeder annotations for this project. This research was funded by generous contributions by philanthropic supporters to the Vermont Center for Ecostudies.

## CONFLICT OF INTEREST

There are no conflicts of interest to be declared.

## AUTHOR CONTRIBUTIONS

D. Narango, P. Sosa-Negrón, and M. Hallworth conceived the ideas; D. Narango developed the methodology, analyzed the data and led the writing of the manuscript; A. Jones, D. Narango, and R. Rebozo collected the data; All authors provided feedback on results and interpretation, contributed critically to the drafts, and gave final approval for publication.

## STATEMENT ON INCLUSION

Our study is a regional synthesis of community-collected data on hummingbirds. As such there was no additional field data collection. Community scientist volunteers from the region were aware of the study’s intentions and outcomes were shared with the community as they developed.

## DATA AVAILABILITY

Data and code to reproduce the analyses will be available from the Dryad Digital Repository upon acceptance of this manuscript.

## Notes

### Competing Interest Statement

The authors have declared no competing interest.

## REFERENCES

Abrahamczyk, S., & Renner, S. S. (2015). The temporal build-up of hummingbird/plant mutualisms in North America and temperate South America. BMC

Andrade, R., Larson, K.L., Franklin, J., Lerman, S.B., Bateman, H.L. and Warren, P.S., 2022. Species traits explain public perceptions of human–bird interactions. Ecological Applications, 32(8), p.e2676.

Archer, C.R., Pirk, C.W.W., Carvalheiro, L.G. and Nicolson, S.W., 2014. Economic and ecological implications of geographic bias in pollinator ecology in the light of pollinator declines. Oikos, 123(4), pp.401–407.

Austin, D.F., 1975. Bird flowers in the eastern United States. Florida Scientist, pp.1–12.

Bates, D., Maechler, M., Bolker, B., Walker, S., Christensen, R.H.B., Singmann, H., Dai, B., Grothendieck, G., Green, P. and Bolker, M.B., 2015. Package ‘lme4’. convergence, 12(1), p.2.

Bertin, R.I., 1982. The ruby-throated hummingbird and its major food plants: ranges, flowering phenology, and migration. Canadian journal of zoology, 60(2), pp.210–219.

Binnie, A., 1965. Some Feeding Habits of the Ruby-Throated Hummingbird. Blue Jay, 23(2).

Brice, A.T., 1992. The essentiality of nectar and arthropods in the diet of the Anna’s hummingbird (Calypte anna). Comparative Biochemistry and Physiology Part A: Physiology, 101(1), pp.151–155.

Carroll, J.E., Marshall, P.M., Mattoon, N.E., Weber, C.A. and Loeb, G.M., 2023. The predation impact of ruby-throated hummingbird, Archilochus colubris, on spotted-wing drosophila, Drosophila suzukii, in raspberry, Rubus idaeus. Crop Protection, 163, p.106116.

Dewitz, J., 2021. National land cover database (NLCD) 2019 products (ver. 3.0, February 2024). US Geological Survey (USGS) Data Release, p.624.

Dick, et al., 2020 Journal of Experimental Biology

Fenster, C. B., Reynolds, R. J., Williams, C. W., Makowsky, R., & Dudash, M. R. (2015). Quantifying hummingbird preference for floral trait combinations: the role of selection on trait interactions in the evolution of pollination syndromes. Evolution, 69(5), 1113–1127.

Fonturbel, F.E., Sepúlveda, I.B., Muschett, G., Carvallo, G.O., Vieli, L. and Murúa, M.M., 2023. Do exotic plants and flower colour facilitate bumblebee invasion? Insights from citizen science data. Flora, 298, p.152200.

Greig, E.I., Wood, E.M. and Bonter, D.N., 2017. Winter range expansion of a hummingbird is associated with urbanization and supplementary feeding. Proceedings of the Royal Society B: Biological Sciences, 284(1852).

Hazlehurst, J.A., Rankin, D.T., Clark, C.J., McFrederick, Q.S. and Wilson-Rankin, E.E., 2021. Macroecological patterns of resource use in resident and migratory hummingbirds. Basic and Applied Ecology, 51, pp.71–82.

Kartesz, J.T., The Biota of North America Program (BONAP), 2015, Taxonomic Data Center, Chapel Hill, N.C. [http://www.bonap.net/tdc]

Lunau, K., Papiorek, S., Eltz, T., & Sazima, M. (2011). Avoidance of achromatic colours by bees provides a private niche for hummingbirds. Journal of Experimental Biology, 214(9), 1607–1612.

Lerman, S.B., Narango, D.L., Andrade, R., Warren, P.S., Grade, A.M. and Straley, K., 2020. Wildlife in the city: human drivers and human consequences. In Urban ecology: its nature and challenges (pp. 37–66). Wallingford UK: CABI.

Lerman, S.B., Larson, K.L., Narango, D.L., Goddard, M.A. and Marra, P.P., 2023. Humanity for habitat: Residential yards as an opportunity for biodiversity conservation. BioScience, 73(9), pp.671–689.

Liaw A, Wiener M (2002). “Classification and Regression by randomForest.” _R News_, *2*(3), 18–22.<https://CRAN.R-project.org/doc/Rnews/>.

Loss, S.R., Will, T. and Marra, P.P., 2015. Direct mortality of birds from anthropogenic causes. Annual Review of Ecology, Evolution, and Systematics, 46(1), pp.99–120.

Marín-Gómez, O.H., Flores, C.R. and del Coro Arizmendi, M., 2022. Assessing ecological interactions in urban areas using citizen science data: Insights from hummingbird–plant meta-networks in a tropical megacity. Urban Forestry & Urban Greening, 74, p.127658.

Maruyama, P.K., Bonizario, C., Marcon, A.P., D’Angelo, G., da Silva, M.M., da Silva Neto, E.N., Oliveira, P.E., Sazima, I., Sazima, M., Vizentin-Bugoni, J. and dos Anjos, L., 2019. Plant-hummingbird interaction networks in urban areas: Generalization and the importance of trees with specialized flowers as a nectar resource for pollinator conservation. Biological conservation, 230, pp.187–194.

Maruyama, P.K., Vizentin_Bugoni, J., Sonne, J., Martin Gonzalez, A.M., Schleuning, M., Araujo, A.C., Baquero, A.C., Cardona, J., Cardona, P., Cotton, P.A. and Kohler, G., 2016. The integration of alien plants in mutualistic plant–hummingbird networks across the Americas: the importance of species traits and insularity. Diversity and Distributions, 22(6), pp.672–681.

Mason, B.M., Mesaglio, T., Barratt Heitmann, J., Chandler, M., Chowdhury, S., Gorta, S.B.Z., Grattarola, F., Groom, Q., Hitchcock, C., Hoskins, L. and Lowe, S.K., 2025. iNaturalist accelerates biodiversity research. BioScience, 75(11), pp.953–965.

Montgomerie, R.D., 1984. Nectar extraction by hummingbirds: response to different floral characters. Oecologia, 63, pp.229–236.

Cubino, J.P., Cavender-Bares, J., Groffman, P.M., Avolio, M.L., Bratt, A.R., Hall, S.J., Larson, K.L., Lerman, S.B., Narango, D.L., Neill, C. and Trammell, T.L., 2020. Taxonomic, phylogenetic, and functional composition and homogenization of residential yard vegetation with contrasting management. Landscape and Urban Planning, 202, p.103877.

Poelen, J.H., Simons, J.D. and Mungall, C.J., 2014. Global biotic interactions: An open infrastructure to share and analyze species-interaction datasets. Ecological informatics, 24, pp.148–159.

Poisot, T., Stouffer, D.B. and Gravel, D., 2015. Beyond species: why ecological interaction networks vary through space and time. Oikos, 124(3), pp.243–251.

Ratto, F., Simmons, B.I., Spake, R., Zamora_Gutierrez, V., MacDonald, M.A., Merriman, J.C., Tremlett, C.J., Poppy, G.M., Peh, K.S.H. and Dicks, L.V., 2018. Global importance of vertebrate pollinators for plant reproductive success: a meta_analysis. Frontiers in Ecology and the Environment, 16(2), pp.82–90.

Rodríguez-Gironés, M.A. and Santamaría, L., 2004. Why are so many bird flowers red?. PLoS biology, 2(10), p.e350.

Rodríguez-Flores CI, Ornelas JF, Wethington S, Arizmendi MdC (2019) Are hummingbirds generalists or specialists? Using network analysis to explore the mechanisms influencing their interaction with nectar resources. PLoS ONE 14(2): e0211855.

Saldivar, J.L.A., Romero, A.N. and Wilson Rankin, E.E., 2022. Community science reveals high diversity of nectaring plants visited by painted lady butterflies (Lepidoptera: Nymphalidae) in California sage scrub. Environmental Entomology, 51(6), pp.1141–1149.

Schuetz, J.G. and Johnston, A., 2019. Characterizing the cultural niches of North American birds. Proceedings of the National Academy of Sciences, 116(22), pp.10868–10873.

Southwick, E. E., & Southwick, A. K. (1980). Energetics of feeding on tree sap by Ruby-throated Hummingbirds in Michigan. American Midland Naturalist, 328–334.

Spence, A.R., Wilson Rankin, E.E. and Tingley, M.W., 2022. DNA metabarcoding reveals broadly overlapping diets in three sympatric North American hummingbirds. The Auk, 139(1), p.ukab074.

Stiles, F.G., 1995. Behavioral, ecological and morphological correlates of foraging for arthropods by the hummingbirds of a tropical wet forest. The Condor, 97(4), pp.853–878.

Tallamy, D.W., Narango, D.L. and Mitchell, A.B., 2021. Do non_native plants contribute to insect declines?. Ecological Entomology, 46(4), pp.729–742.

Weidensaul, S., T. R. Robinson, R. R. Sargent, M. B. Sargent, and T. J. Zenzal (2020). Ruby-throated Hummingbird (Archilochus colubris), version 1.0. In Birds of the World (P. G. Rodewald, Editor). Cornell Lab of Ornithology, Ithaca, NY, USA. 10.2173/bow.rthhum.01

Wolf, N., Smeltz, T.S., Cook, C. and Martinez del Rio, C., 2023. Using stable isotopes in hummingbird breath to estimate reliance on supplemental feeders. Ecology and Evolution, 13(2), p.e9799.

Zenzal, T.J. and Moore, F.R., 2019. Resource use and defence by ruby-throated hummingbirds during stopover. Behaviour, 156(2), pp.131–153.

